# Hybrid Songbirds are Deficient in Learning and Memory

**DOI:** 10.1101/227298

**Authors:** Michael A. McQuillan, Timothy C. Roth, Alex V. Huynh, Amber M. Rice

## Abstract

Identifying the phenotypes underlying postzygotic reproductive isolation is crucial for fully understanding the evolution and maintenance of species. One potential postzygotic isolating barrier that has not yet been examined is learning and memory ability in hybrids. Learning and memory are important fitness-related traits, especially in scatter-hoarding species, where accurate retrieval of hoarded food is vital for winter survival. Here, we test the hypothesis that learning and memory ability can act as a postzygotic isolating barrier by comparing these traits among two scatter-hoarding songbird species, black-capped (*Poecile atricapillus*), Carolina chickadees (*Poecile carolinensis*), and their naturally occurring hybrids. In an outdoor aviary setting, we find that hybrid chickadees perform significantly worse on an associative learning spatial task and are worse at solving a novel problem compared to both parental species. Deficiencies in learning and memory abilities could therefore contribute to postzygotic reproductive isolation between chickadee species. Given the importance of learning and memory for fitness, our results suggest that these traits may play an important, but as yet overlooked, role in postzygotic reproductive isolation.

## Introduction

A fundamental goal in evolutionary biology is to understand the origin and maintenance of species. Natural hybridization – when distinct species mate and produce offspring of mixed ancestry – provides a window into the process of species formation and can highlight the barriers that reduce gene flow between species (Abbott et al. 2013). For example, hybridization often results in the production of unfit hybrid offspring that are selected against (Coyne and Orr 2004). This selection against hybrids, or postzygotic reproductive isolation, can maintain the integrity of species boundaries despite ongoing gene flow (Servedio and Noor 2003). Alternatively, hybridization can cause two species to fuse back into one as speciation moves in reverse (Taylor et al. 2006). Finally, hybridization can stimulate evolutionary novelty through the formation of completely new hybrid lineages (Gompert et al. 2006; Mallet 2007; Trier et al. 2014), or by promoting the transfer of adaptive genetic material across species boundaries (Arnold 1997; Grant and Grant 2010). Although the evolutionary outcomes of hybridization may vary widely, they all influence the speciation process in some way. Therefore, identifying the mechanisms underlying selection against hybrids and postzygotic isolation is crucial for fully understanding how new species are formed and species barriers are maintained.

Two important fitness-related traits that have not been examined as potential postzygotic isolating barriers are learning and memory. Learning has a well-established role in *prezygotic* reproductive isolation, however (Verzijden et al. 2012; Dukas 2013). For example, many species learn their mate preferences through sexual imprinting, where juveniles learn the phenotypes of their parents and later select mates based on those phenotypes (Cate and Vos 1999; Verzijden et al. 2008; Kozak et al. 2011). Species can also learn the traits that are themselves the targets of mate choice, such as birdsong (Sorenson et al. 2003; Beecher and Brenowitz 2005). Despite a demonstrated role for learning in prezygotic isolation, a potential role in postzygotic isolation has not been adequately tested. A small number of studies have examined learning and memory abilities in hybrids (Proops et al. 2009; Osthaus et al. 2013; Hoedjes et al. 2014), and in some cases found that hybrids exhibited enhanced cognition compared to parental species (e.g. Proops et al. 2009; Osthaus et al. 2013). However, these studies all examined cognition in hybrids created by crossing domesticated species or inbred lines. In such cases, hybrid cognitive performance may reflect enhanced heterozygosity and relief from inbreeding depression. Therefore, it is difficult to generalize these results to cases of natural hybridization. Thus, the potential role of learning and memory in postzygotic isolation remains relatively unexplored. Here, our goal was to fill this gap by assessing learning and memory abilities in two songbird species and their naturally occurring hybrids, for which learning and memory traits are important for survival.

Learning and memory are important for fitness and survival in many species (e.g. Hollis et al. 1997; Dukas and Bernays 2000; Maille and Schradin 2016), and this has been particularly well demonstrated in species that are ‘scatter-hoarders’ (Vander Wall 1990; Shettleworth 1998). Scatter-hoarding species cache (store) food items in hundreds to thousands of locations scattered around their habitat. Individuals must then rely on cognitive traits, like learning and memory, to accurately retrieve cached food at a later time (Pravosudov and Smulders 2010). The ability to accurately retrieve caches is particularly important for winter survival, especially in harsh environments where alternate sources of food are scarce or unpredictable (Pravosudov and Clayton 2002). Furthermore, learning and memory traits in scatter-hoarders are both heritable and variable among species, setting the stage for possible hybrid breakdown in these traits upon interspecific hybridization (Brodbeck 1994; Roth et al. 2010, 2012; Pravosudov and Roth 2013).

Our two study species, the black-capped (*Poecile atricapillus*) and Carolina (*P. carolinensis*) chickadee, are well-studied scatter-hoarding passerines that hybridize extensively along a narrow hybrid zone stretching from New Jersey to Kansas (Bronson et al. 2005; Curry 2005; Taylor et al. 2014a; McQuillan and Rice 2015; **Fig. 1a**). These species are sister taxa (Harris et al. 2014), and display ~6% mitochondrial DNA sequence divergence (Gill et al. 2005). While selection against hybrid chickadees is strong at early life stages due to reduced hatching success (Bronson et al. 2003, 2005), little is known about additional selection pressures on adult hybrids (but see Olson et al. 2010).

**Figure 1.**
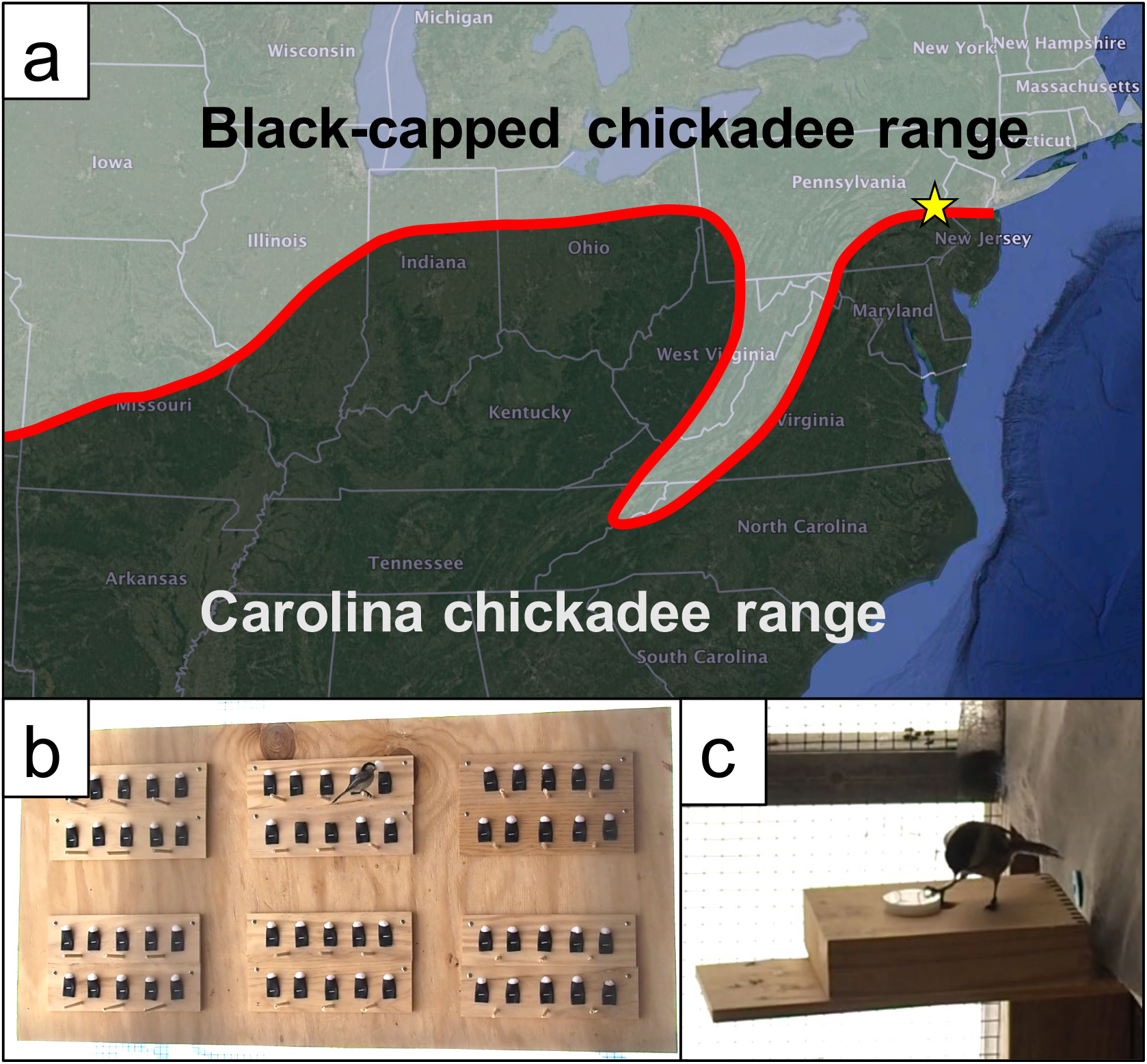
(A) Approximate location of black-capped and Carolina chickadee hybrid zone shown in red. Sampling location for this study in eastern Pennsylvania is shown as yellow star. (B) Chickadee interacting with ‘caching array’ used in associative learning spatial task. (C) Chickadee performing novel problem-solving test.

Learning and memory abilities in chickadees are heritable and shaped by natural selection. For example, black-capped chickadees inhabiting harsh winter environments, where selection is expected to favor enhanced cognition, cache more food and display significantly better spatial learning, memory, and problem-solving abilities than birds from mild environments (Clayton and Pravosudov 2002; Roth et al. 2010, 2012). In addition, chickadees from these harsh environments possess significantly more hippocampal neurons (the area of the brain associated with spatial memory), and display higher levels of hippocampal neurogenesis than mild-environment birds (Roth and Pravosudov 2009; Roth et al. 2011). These behavioral and hippocampal differences remain when birds from the divergent environments are raised under common laboratory conditions, suggesting a heritable genetic basis (Roth et al. 2010, 2012). Carolina chickadees are also prodigious scatter-hoarders (Lucas and Zielinski 1998; Lucas et al. 2006), and one might expect learning and memory to be important traits for survival, especially at the northern portion of their range where winters can be harsh and hybridization occurs (**Fig. 1a**).

In this study, we test the hypothesis that learning and memory have the potential to function in postzygotic reproductive isolation by comparing the learning and memory abilities of wild-caught black-capped, Carolina, and hybrid chickadees from a hybrid zone population. Specifically, we measure individual performance on an associative learning spatial task and a novel problem-solving test. If hybrids perform worse in these tests of learning and memory compared to one or both parental species, then these traits may represent a novel postzygotic isolating barrier. In contrast, if hybrids display equal or superior abilities compared to parental species, then learning and memory are unlikely to act as postzygotic barriers in this system, and may even provide adaptive benefits to hybrids.

## Materials and Methods

### Field Methods and Aviary Housing

We caught black-capped, Carolina, and hybrid chickadees (total n=50) using mist nets at bird feeders or using song playback over the course of 13 months (Jan 2016–Feb 2017), excluding most of the breeding season (April-May). We caught birds at two geographically proximate sites within the hybrid zone in eastern Pennsylvania (20 km apart; Lehigh University 40°36’5.19”N, 75°21’34.08”W; Jacobsburg State Park 40°47’3.97”N, 75°17’34.67”W) (**Fig. 1a**). Upon capture, we banded, measured, and weighed each bird, and took a blood sample for ancestry (McQuillan et al. 2017) and sex determination (Griffiths et al. 1998). We then transported each bird by car to an outdoor aviary at Lehigh University, where birds were housed individually in aviary compartments measuring 1.5 × 2 × 2.5 m. Birds were visually, but not aurally isolated from each other. We covered the outside of the aviary with house wrap to prevent birds from seeing out, while still allowing ambient light to pass through. All behavioral testing took place in home aviary compartments. We provided birds with *ad libitum* sunflower seeds, pine nuts, and vitamin-supplemented water. We used wax worms, a highly desirable food item, in behavioral trials (see below). We outfitted each aviary compartment with 10 small rubber caching pockets (each 2.5 × 4 cm) that were accessible by perches. We did this so that birds would acclimate to caching and retrieving food from these pockets, which were similar to those used in the associative learning spatial task, described below. After behavioral testing was complete (birds spent an average of 3 weeks in captivity), we released birds at the point of capture. All procedures were approved by Lehigh University’s Institutional Animal Care and Use Committee (Protocol #175).

### Genetic Determination of Species Ancestry

Because black-capped, Carolina, and hybrid chickadees are morphologically similar and song is not a reliable species-identifier within the hybrid zone (Kroodsma et al. 1995), we utilized genetic tools to assign ancestry to each bird. Briefly, we genotyped all birds at 10 species-diagnostic single nucleotide polymorphism (SNP) markers (McQuillan et al. 2017). Because chickadees in our population are highly admixed and include pure-species, F1, as well as advanced-generation and backcrossed hybrids (McQuillan et al. 2017), we used STRUCTURE (Hubisz et al. 2009) to estimate admixture proportions and assign ancestry categories for each bird. To do this, we combined the genotypes of our test subjects with a larger dataset containing genotypes of individuals (N=301) from multiple Pennsylvania hybrid-zone populations, as well known pure-species individuals from allopatric populations of both species (New York and Louisiana, USA). We ran STRUCTURE on this larger dataset, using the same program settings as those used in McQuillan et al. 2017 (**Fig. S1**). Birds with admixture values that fell within the average 90% credible interval of the known pure individuals in the dataset were classified as either a pure black-capped or Carolina individual. In contrast, birds with admixture values outside of the average 90% credible interval for the known parentals were considered hybrids.

### Associative Learning Spatial Task

We subjected birds to an associative learning spatial task (following Roth et al. 2012). We used a large plywood ‘caching array,’ measuring 1.25 × 0.6 m with 60 rubber pockets identical to those used during acclimation, and accessible by perches. The contents of each pocket could be concealed by placing a white, craft ‘pom-pom’ ball over the opening, which birds had to remove in order to investigate the pocket’s contents (**Fig. 1b**). All birds were able to remove the balls from the pockets. On the first day of the test, we placed a wax worm in one of the 60 pockets (chosen randomly for each bird), covered it and all other pockets with the balls, introduced the array to the bird’s aviary compartment, and allowed the bird to remove the pom-pom balls and find the worm. For the next 9 days, once per day, we presented the same caching array to the bird with all pockets covered and a wax worm concealed in the same pocket. Care was taken to hang the caching array on the aviary wall in the same place each day. Each day, we recorded the number of inspections (defined as the number of pom-pom balls removed) required to successfully locate the worm. We tested 11 black-capped, 8 Carolina, and 10 hybrid chickadees on this task.

Because some of the birds we tested had been used previously to collect pilot data for another experiment using the caching array (total n=10; 2 black-capped, 5 Carolina, 3 hybrid), we tested for an effect of this prior experience on performance on the associative learning spatial task. We found a marginally non-significant interaction between this prior experience and testing day (GLMM χ^2^(1)=3.815, p=0.051). Although not statistically significant, we sought to eliminate any potential influence of prior experience on our results by limiting our subsequent analysis to days 4 through 10 (hereafter referred to as testing days 1 through 7). During this time window, the performance of birds with and without prior caching array experience converged (**Fig. S2**; GLMM; prior experience × testing day, χ^2^(1)=0.370, p=0.543; prior experience main effect, χ^2^(1)=1.692, p=0.193). Importantly, limiting our analysis to this time window did not qualitatively change our results.

### Novel Problem-Solving Test

To evaluate each bird’s innovativeness and general learning ability, we assessed each bird’s ability to solve a reward-based, novel problem (following Roth et al. 2010). The problem-solving test required birds to physically move a circular nylon washer with a transparent coating (3.3 cm diameter, 6 g, ~1/2 the mass of the bird) from covering a 1.75 cm well in a wooden block (15 × 10 × 4 cm) that contained a waxworm (**Fig. 1c**). The test was designed so that birds could see through the circular washer and recognize that a food reward was concealed underneath, but the reward could only be retrieved by sliding the washer to the side. On the day of the problem-solving test, we deprived birds of food for one hour in the morning, after which we introduced the problem-solving apparatus to the bird’s home compartment. We recorded whether or not the bird was able to solve the problem (by moving the washer and recovering the worm) in a one-hour maximum time-frame. We conducted a second test in the afternoon, with the two tests spaced by at least 4 hours. We tested 14 black-capped, 6 Carolina, and 16 hybrid chickadees (total n=36) on this problem-solving test. Included in this group of 36 birds were 15 birds that had previously performed the associative learning spatial task.

All birds were habituated to finding wax worms in the well of the problem-solving apparatus and to the presence of the nylon washer, which was mounted adjacent to the well during habituation (>40 hours of habituation time per bird). To control for motivation and the birds’ willingness to feed from the problem-solving apparatus, we conducted a pre-trial control in the afternoon of the day preceding the problem-solving test. For the pre-trial, we introduced the problem-solving apparatus to the bird with an un-concealed wax worm and the nylon washer mounted adjacent to the well. We measured the latency in seconds for the bird to land on the apparatus and take the worm. Birds were also food deprived for one hour prior to this pre-trial control.

### Statistical Analysis

To test for performance differences among ancestry groups in the associative learning spatial task, we fit generalized linear mixed models (GLMMs) using maximum likelihood to our data. We specified log link functions in our models and used the ‘lme4’ package in R, version 3.4.1 (R Core Team, 2017; Bates et al. 2015). Specifically, we fit GLMMs with ‘score’ (the number of pom-pom balls removed before the bird recovered the worm) as our response variable. We included the following independent variables as fixed effects, including all possible interactions: Ancestry (black-capped, Carolina, or hybrid), sex, testing day, and season (spring, fall, or winter). In order to account for the repeated-measures nature of our dataset (i.e. we measured each bird’s performance once per day for multiple days), we specified a random intercept, as well as a random slope for each bird across testing days in our model. Once we had constructed this ‘full’ model, we performed a step-wise model simplification procedure by removing the least significant variable, starting with the highest order interactions. After variable removal, we performed likelihood ratio tests (LRT) to compare models with and without each focal term. If the simplified model explained significantly less variation in the response variable, then the focal term was retained. We continued this process until we were left with a ‘best-fit’ model, which contained only those fixed effects that were significant predictors, according to the likelihood ratio tests. If a variable was found to be a significant predictor, we assessed post-hoc pairwise contrasts of the levels within that factor using least-square means (LSM) in the R package ‘lsmeans’ (v. 2.27-2; Lenth 2016).

To test for differences among the ancestry groups in ability to solve the novel reward-based problem, we compared the proportion of individuals from each ancestry group that were able to successfully solve the problem at least once (of two total trials), using Fisher’s exact tests.

## Results

For the associative learning spatial task, hybrids performed worse across the testing period than either pure species. Our best-fit GLMM was significant overall (LRT compared to null model, χ^2^(4)=31.10, p<0.001), and contained ancestry (χ^2^(2) = 8.11, p=0.017), sex (χ^2^(1)=8.32, p=0.004), and testing day (χ^2^(1)=17.62, p<0.001) as fixed effects. Our post-hoc pairwise comparisons indicated that hybrids required more inspections to recover the wax worm across the testing period than pure black-capped (estimated LSM contrast=0.437, p=0.033) and Carolina (LSM contrast=0.525, p=0.018) birds (**Fig. 2**). There was no significant difference in performance between the two pure-species groups (LSM contrast= 0.088, p=0.890). Additionally, male birds made significantly fewer inspections to recover the wax worm, on average, than female birds (LSM contrast= 0.503, p=0.002). The significant effect of testing day on test performance indicated that learning occurred across all ancestry groups, with birds requiring fewer inspections to recover the food reward as the test progressed. Interestingly, female hybrids were the worst performers on the associative learning spatial task (**Fig. 3**). Although this sex × ancestry interaction was not retained as a significant predictor in the best-fit model, likely due to low power, it suggests the intriguing possibility that hybrid cognition may follow Haldane’s Rule (see discussion).

**Figure 2.**
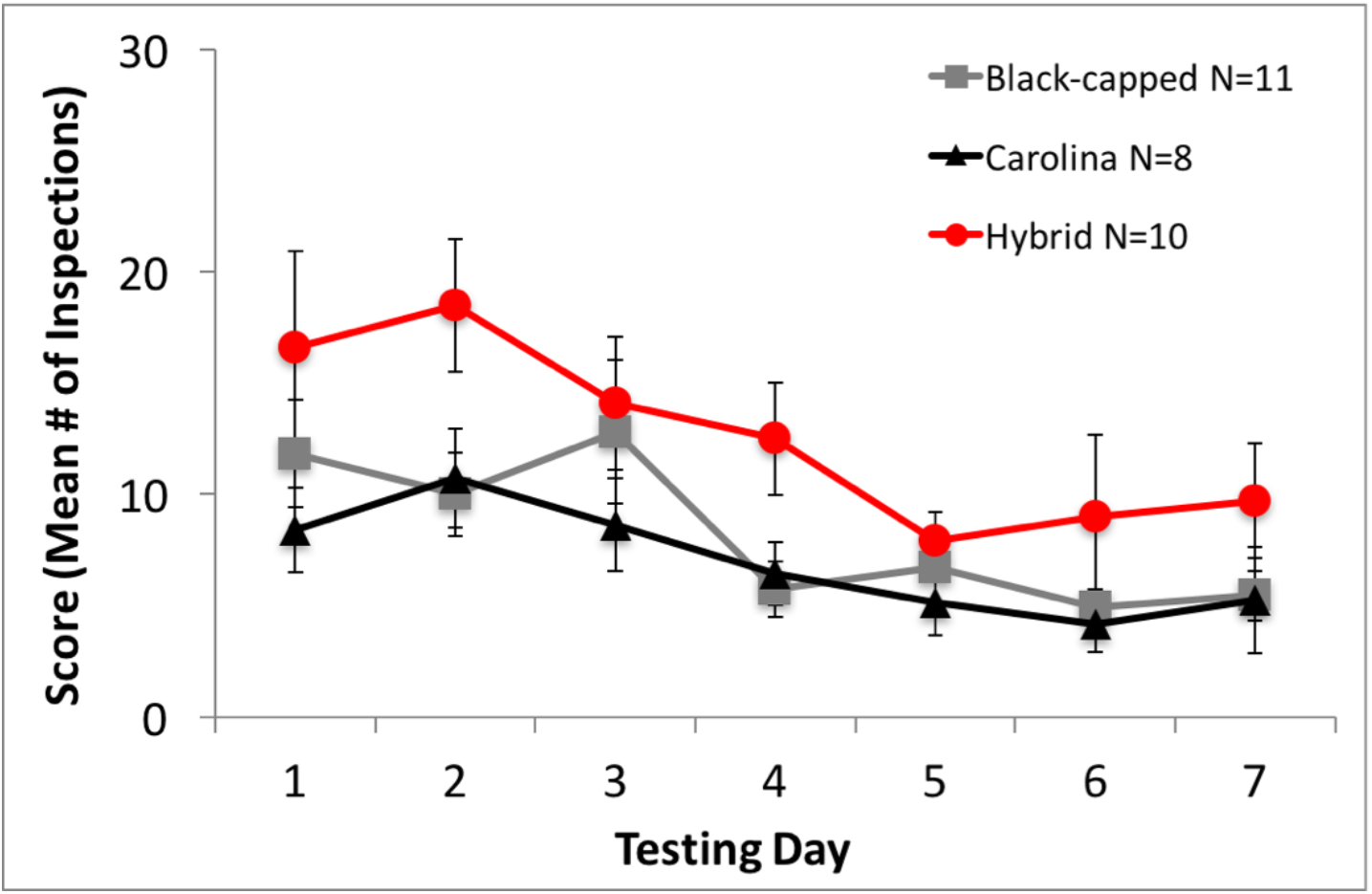
Associative learning spatial task results, showing performance over the testing period for the three ancestry groups. Hybrid chickadees require significantly more inspections, on average, than both pure-species groups. Data points denote mean ± SEM.

**Figure 3.**
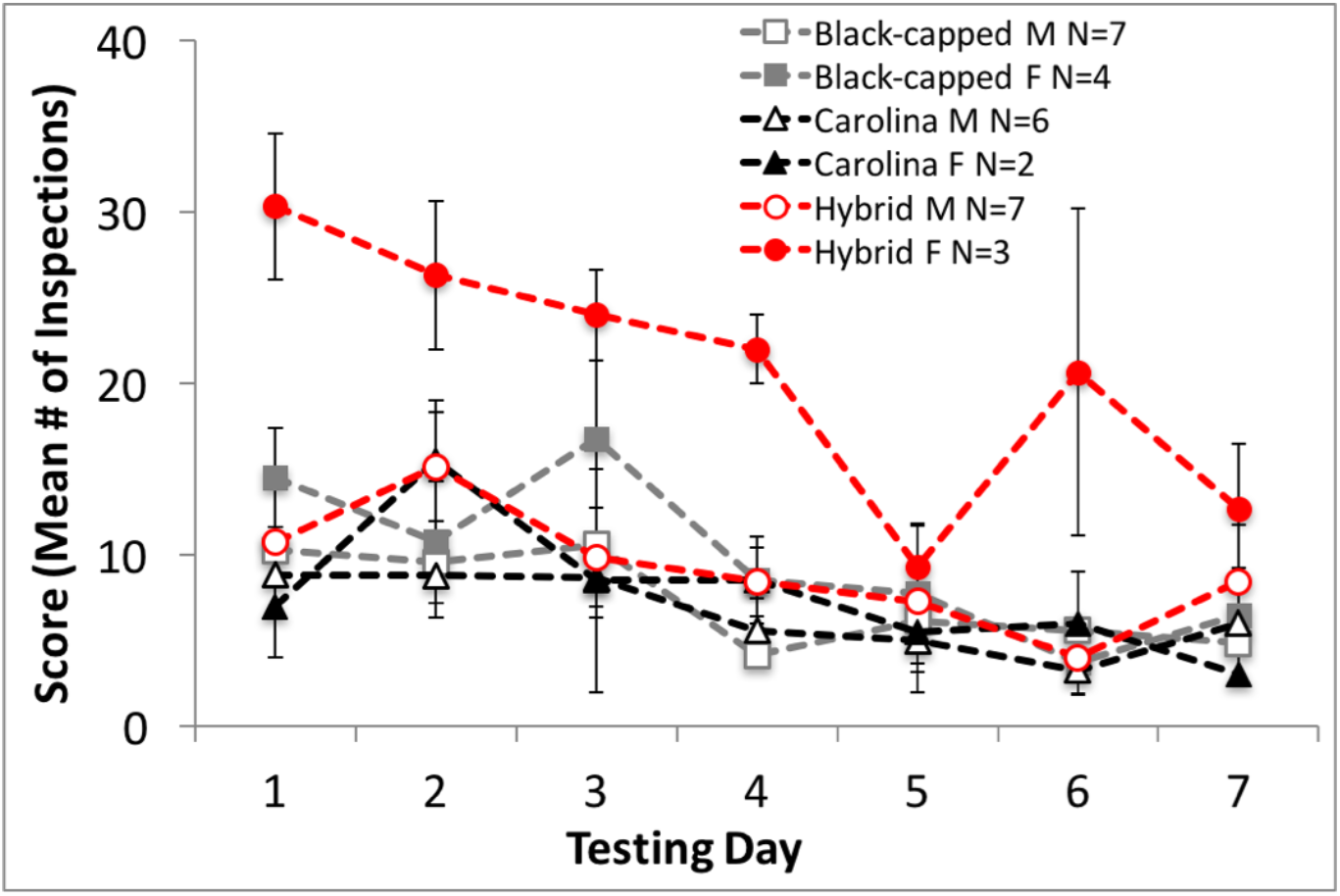
Performance on the associative learning spatial task split by ancestry and sex. Hybrid females are the worst performers on associative learning spatial task (red dashed line, closed circles). Data points denote mean ± SEM.

Hybrids were also worse at problem solving. In the novel problem-solving test, 13/14 (93%) black-capped, 6/6 Carolina (100%), and 10/16 (62.5%) hybrids were able to solve the problem at least once (**Fig. 4a**). These differences in success were marginally non-significant (Fisher’s exact test, p=0.054). However, the lack of statistical significance may be due to a lack of power from the relatively small Carolina chickadee sample size. Because we were most interested in testing hybrid cognitive abilities relative to pure species, and because the black-capped and Carolina chickadees exhibited similar rates of success (**Fig. 4a**), we combined black-capped and Carolina birds into a single, pure-species category and compared it to the hybrid group. When analyzed in this way, hybrids were significantly less likely to solve the problem than pure-species individuals across the two trials (**Fig. 4b**; p=0.0298, Fisher’s exact test; 62.5% vs. 95% success, respectively). All birds that solved the problem during the first trial also successfully solved in the second trial. There were no significant differences between ancestry groups in the latency to take the worm in the pre-trial control, indicating that all birds were equally motivated to feed from the problem-solving apparatus (Kruskal-Wallis χ^2^(2)=2.78, p=0.248). Additionally, differences among the ancestry groups in problem-solving ability cannot be explained by differences in body mass (ANOVA F(2,32)=1.469, p=0.245; **Fig. S3**).

**Figure 4.**
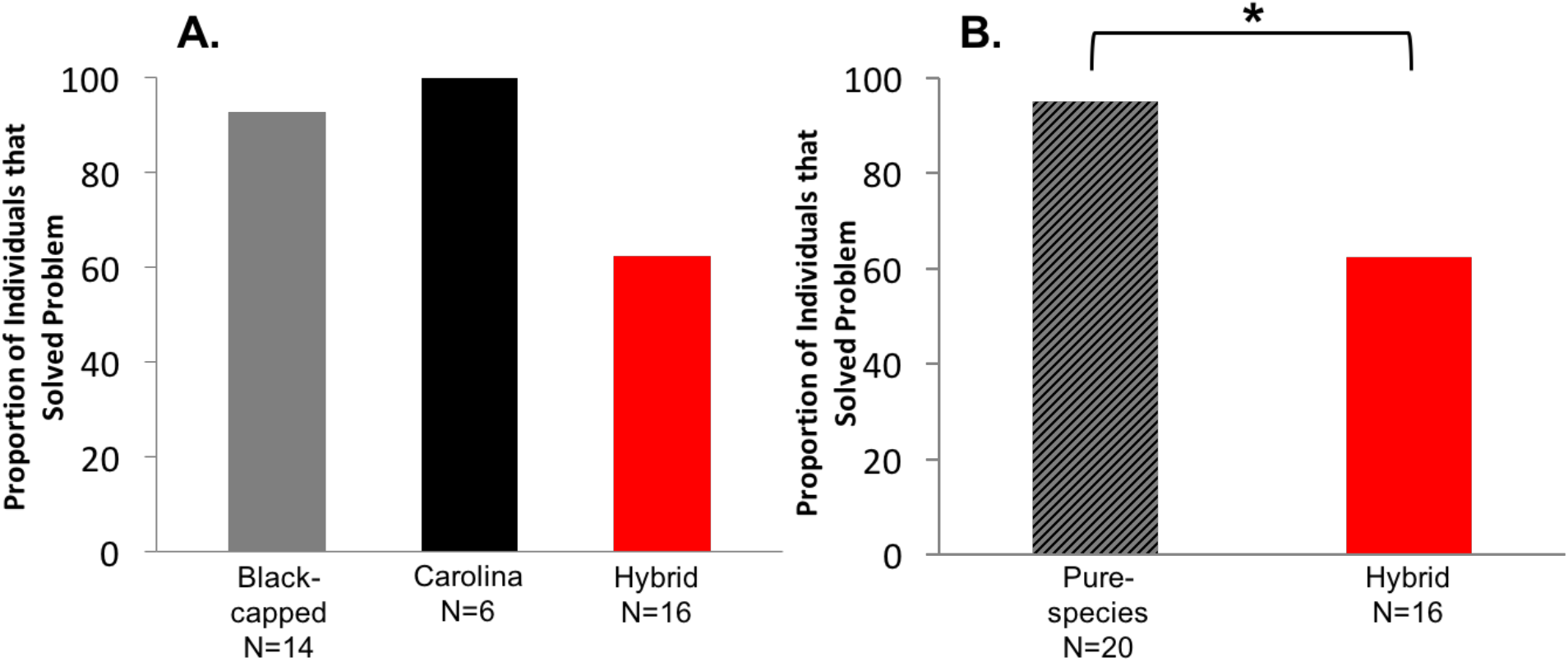
Novel problem-solving results. (A) Proportion of individuals from each ancestry group that were able to solve the problem at least once (of two total trials). Hybrids show a non-significant trend towards being less likely to solve the problem (Fisher’s exact test, p=0.054). (B) Hybrids are significantly less likely to solve the problem compared to a combined pure-species group (Fisher’s exact test, p=0.0298).

## Discussion

Overall, our results suggest that hybrid chickadees are deficient in learning and memory traits relative to their pure-species counterparts. Hybrid birds performed worse on average than pure-species individuals at an associative learning spatial task (**Figs. 2,3**), and were less likely to solve a reward-based novel problem than pure-species birds (**Fig. 4**). These results suggest a role for learning and memory in postzygotic isolation, which has not been examined previously. Our behavioral results likely have real-world implications for scatter-hoarding chickadees in nature, as learning and memory are important fitness-related traits in these species.

Accurate retreival of cached food is crucial for winter survivial, and natural selection shapes the cognitive traits necessary to achieve this (Pravosudov and Roth 2013). Previous work uncovered heritable variation in caching-related learning and memory traits between populations of black-capped chickadees from different environments (Pravosudov and Clayton 2002; Roth et al. 2010, 2012). These results suggest that learning and memory traits in chickadees are important for survivial, potentially under genetic control, and shaped by natural selection. We extended this line of reasoning by examining the learning and memory abilites of hybrid chickadees, and comparing them against pure-species individuals from the same environment. The fact that hybrid chickadees display inferior learning and memory abilities relative to pure-species birds suggests that hybrids may suffer a fitness disadvantage in nature, particularly in their ability to accurately retreive cached food. There are multiple possible explanations for this deficiency in hybrids.

One possible explanation for hybrid deficits in our cognitive tests is the accumulation of genetic incompatibilities among loci underlying learning and memory traits in hybrid genomes. Genetic incompatibilites arise when alleles at two or more loci diverge in geographically isolated species, and recombine in hybrid genomes upon secondary contact (Dobzhansky 1936; Muller 1942). If genes underlying learning and memory traits have diverged in black-capped and Carolina chickadees, negative genetic interactions between these loci could occur in hybrids, leading to breakdown in these traits. One ubiquitous outcome of hybrid genetic incompatibilities is Haldane’s Rule, which states that within hybrid offspring, the heterogametic sex (i.e. the sex possessing two different sex chromosomes) is more likely to be absent, rare, or sterile (Haldane 1922). This rule applies whether males are heterogametic, as in mammals and *Drosophila*, or whether females are heterogametic, as in birds and Lepidoptera. Haldane’s Rule has been upheld in virtually all taxa that have been surveyed (Orr 1997; Delph and Demuth 2016). Of the multiple explanations for Haldane’s Rule, all are genetic in nature. The explanation with the most support, termed ‘dominance theory’ (Turelli and Orr 1995), posits that the heterogametic sex (females in birds) will be negatively affected by any and all incompatibility loci located on the sex chromosmes, regardless of whether the incompatible alleles act dominantly or recessively. In contrast, the homogametic sex (males in birds) will only be negatively affected by the subset of sex-linked incompatibility loci that act in a relatively dominant fashion.

Consistent with Haldane’s Rule, our results on the associative learning spatial task suggest that female hybrids have the worst memories (**Fig. 3**). If hybrid females are less able to accurately retrieve cached food in nature, then their ability to survive particularly harsh winter conditions may be reduced. Interestingly, Bronson et al. (2005) found a distinct paucity of hybrid female chickadees along a hybrid zone transect in Ohio. The authors concluded that female hybrids may have reduced viability. Our results provide a potential explanation for this result. Furthermore, a genomic analysis of introgression patterns across the chickadee hybrid zone in Pennsylvania found that a majority of loci putatively underlying reproductive isolation are located on the Z chromosome (in birds, females are heterogametic ZW, while males are homogametic ZZ; Taylor et al. 2014b). This result would predict that female hybrid chickadees are likely to experience greater negative effects from sex-linked genetic incompatibility loci than male hybrids. In contrast, we did not discover an effect of sex in the novel problem solving test: Of the six hybrids that were unable to solve the problem, three were male and three were female.

Another possible explanation for our results is that hybrid chickadees are not deficient in their learning and memory abilities, but rather are deficient in some other respect, such as their general health or body condition. In other words, worse performance on our cognitive tests may be an artifact of reduced overall hybrid quality. For example, Turissini et al. (2017) found that lab-generated *Drosophila* hybrids were less capable of finding food than their pure-species parents. If hybrid chickadees are also worse foragers than pure-speices individuals, impaired cognitive performance may simply result from the fact that hybrids have a lower nutritional state. In rats, nutritional stress leads to anatomical defects in the hippocampus, which is associated with reduced performance on spatial memory tasks (Jordan et al. 1981; Levitsky and Strupp 1995). However, we see no evidence that hybrid chickadees are of lower body condition than pure-species birds in our hybrid zone transect as a whole, or in the subset of birds used for this study (**Fig. S4**).

Related to this point, it is also possible that deficient learning and memory in hybrids could result from overall metabolic dysfunction. Because the brain is a metabolically costly organ, breakdown in hybrid metabolism could result in an associated breakdown in learning and memory. In an Ohio hybrid zone transect, hybrid chickadees display higher mass-corrected basal metabolic rates compared to both parental species, suggesting less efficient hybrid metabolism (Olson et al. 2010). Similar results have been found in hybrid flycatchers (Mcfarlane et al. 2016). Future work should aim to disentangle the possible consequences of metabolic breakdown on learning and memory from other neurological causes for breakdown in these traits. More generally, future work should examine the relative contributions of genetic/epigenetic factors versus plastic responses to the environment on hybrid cognition. For example, hybrids may experience the environment differently than pure-species individuals, and this difference in life experience could affect cognitive performance. This could be controlled for by hand-rearing pure-species and hybrid individuals under identical laboratory conditions and subjecting them to learning and memory tests as adults (as in Roth et al. 2010, 2012).

The fact that hybrids are less able to solve a novel problem than pure-species birds (**Fig. 4**) also likely has real-world implications in this system. In unpredictable environments, animals must either invent new behaviors, or flexibly adjust established behaviors in order to solve novel problems (Reader and Laland 2003). The ability to solve novel problems is often used as a measure of an animal’s ability to innovate (e.g. Morand-Ferron et al. 2011; Cauchard et al. 2013). Environmental perturbations as a result of climate change will add to the unpredictability of many environments, perhaps increasing selective pressure for innovativeness and problem-solving abilities. If hybrids are less able to solve problems and innovate than their pure-species counterparts, this may represent a selective disadvantage in nature. As the frequency of hybridization is expected to rise under climate change (Chunco 2014), future work should examine hybrid problem-solving abilities in other systems inhabiting unpredictable environments.

Identifying the barriers that prevent gene flow between closely related species is crucial in order to understand how new species arise. Much work on postztgotic isolation has focused on identifying the genetic underpinnings of hybrid sterility and inviability, often in model laboratory systems. However, less attention has been paid to maladaptive behaviors in hybrids that potentially contribute to reproductive isolation (but see Noor 1997a; Delmore and Irwin 2014; Schmidt and Pfennig 2016). Here, we have shown evidence that naturally occurring hybrids are deficient in learning and memory abilities relative to their pure-species parents. This may be an important and widespread, but as yet overlooked, source of postzygotic reproductive isolation. One additional goal of future studies should be to evaluate the degree to which hybrids display these deficiencies in their natural environments (sensu Croston et al. 2016). Given the widespread nature of hybridization (Mallet 2005), examining potential behavioral sources of postzygotic isolation is paramount for a complete understanding of the speciation process.

## Acknowledgements

We would like to acknowledge Jessie Brill, Joseph Skibbens, and Ryan Herbert for assistance in the field and for helpful discussion. We also thank Richard Wiltraut and Robert Neitz for assistance at our Jacobsburg State Park field site. This work was funded by a Sigma Xi Grant-in-Aid of Research (MAM), the George W. Barlow Award from the Animal Behavior Society (MAM), the Gordon C. Thorne graduate fellowship from Lehigh University (MAM), and a Lehigh University Faculty Innovation Grant (AMR).

